# Gender differences in submission behavior exacerbate publication disparities in elite journals

**DOI:** 10.1101/2023.08.21.554192

**Authors:** Chaoqun Ni, Isabel Basson, Giovanna Badia, Nathalie Tufenkji, Cassidy R. Sugimoto, Vincent Larivière

## Abstract

Women are particularly underrepresented as leading authors of papers in journals of the highest impact factor, with substantial consequences for their careers. While a large body of research has focused on the outcome and the process of peer review, fewer articles have explicitly focused on gendered submission behavior and the explanations for these differences. In our study of nearly five thousand active authors, we find that women are less likely to report having submitted papers to journals of the highest impact (e.g., *Science, Nature*, or *PNAS*) andto submit fewer manuscripts, on average, than men when they do submit. Women were more likely to indicate that they did not submit their papers (in general and their subsequently most cited papers) to high-impact journals because they were advised not to. In the aggregate, no statistically significant difference was observed between men and women in how they rated the quality of their work. Nevertheless, regardless of discipline, women were more likely than men to indicate that their “*work was not ground-breaking or sufficiently novel”* as a rationale for not submitting to one of the listed prestigious journals. Men were more likely than women to indicate that the “*work would fit better in a more specialized journal*.*”* We discuss the implications of these findings and interventions that can serve to mitigate the disparities caused by gendered differences in submission behavior.

**Significance:** Publishing in high-impact scholarly journals has a significant effect on researchers’ careers. Our findings identify factors that affect submission to *Science, Nature*, and the *Proceedings of the National Academy of Sciences of the United States of America* (*PNAS*) and explore whether there is a relationship between gender and desk rejections or submission rates. We found no relationship between gender and reported desk rejection and a relationship between gender and reported submissions, with men having a greater number of submissions. Women were more likely than men to indicate that their “*work was not ground-breaking or sufficiently novel”* for the listed prestigious journals and that they were advised against submitting to these venues. Men were more likely to indicate that the “*work would fit better in a more specialized journal*.*”*

## Introduction

The rise of the research evaluation system has created a market of intense competition for a few hallowed venues. Scientific capital is strongly concentrated in generalist journals founded in the late nineteenth and early twentieth Centuries (Baldwin, 2015), such as *Science* (1880), *Nature* (1869), and *Proceedings of the National Academy of Sciences of the United States of America* (PNAS, 1914). Articles published in these journals garner greater news media attention, and authors tend to receive more career opportunities and grants (Reich, 2013). Since publishing in highly influential journals leads to wider dissemination and visibility of researchers’ work, the volume of manuscript submissions to these journals is high and acceptance low, with a large proportion of manuscripts rejected by editors without undergoing peer review. It is noted on *Science’s* author portal that less than 6% of originally submitted papers are accepted (Science, 2025). *Nature* also notes low acceptance rates on its website, from 8-12% depending on the year, and states that “*many submissions are declined without being sent for review*” (Nature, 2025). A similar practice is espoused by the *Proceedings of the National Academy of Sciences of the United States of America (PNAS)*, with most publications rejected before peer review and a final acceptance rate of 14.7% (PNAS, 2025). Prestige and hierarchy among journals are not inherently negative for scholarly communication; however, if entry into these highly visible venues varies for different sociodemographic populations of authors, it may have adverse consequences for science (Kozlowski et al., 2022).

Editors and peer reviewers evaluate both the quality of manuscripts submitted as well as whether the topic fits within the scope of the journal. Idealized gatekeeping reflects the Mertonian norms of organized skepticism and universalism, evaluating scientific work independently of the social characteristics of authors (Merton, 1968). However, various critiques of scholarly peer review have been discussed (Ware, 2008), among which is a concern of social biases, that is, “differential evaluation of an author’s submission as a result of her/his perceived membership in a particular social category” (Lee et al., 2013). These biases are not necessarily conscious; yet, if no biases were present in the case of peer review, then “we should expect the rate with which members of less powerful social groups enjoy successful peer review outcomes to be proportionate to their representation in *submission rates*” (Lee et al., 2013). Results on gender disparities and bias in scholarly publishing have arrived at mixed results (Borsuk et al., 2009; Day et al., 2020; Djupe et al., 2019; Edwards et al., 2018; Fox & Paine, 2019; Gilbert et al., 1994; Grossman, 2020; Helmer et al., 2017; Murray et al., 2018; Squazzoni et al., 2021; Walker et al., n.d.). However, many of these fail to disentangle components of the publication process, focusing only on the outcome of peer-reviewed publications rather than the effect of different groups having different submission rates or desk rejections. These components have fundamentally different attributes—with one being an effect of self-selection and the other a potential indicator of bias. We emphasize here that observations of disparities are a necessary, but insufficient indicator of bias as there are several other potential explanations for disparities. We focus, therefore, on the concept of disparity.

Most studies focused on disparities in science publishing have demonstrated that women submit fewer papers than men, with little difference in desk rejections (or a disparity in favor of women) (Bendels et al., 2018; Garand & Harman, 2021; Grossman, 2020; Martinsen et al., 2022; Squazzoni et al., 2021). The lower submission rate is particularly pronounced in the most prestigious journals (Bendels et al., 2018). Interviews with women authors explained these lower submission rates with higher selectivity: they argued that they only submit when they are confident the paper would be accepted, not wanting to risk a rejection as they have limited time for conducting research and attending to lengthy reviews (Closa et al., 2020). Women were also less likely than men to indicate that they submit their manuscripts to top journals in their field as their first choice and more likely than men to select a journal based on whether a journal is most likely to accept the manuscript (Djupe et al., 2019). Other perceptions and behavioral differences have also been postulated to account for the differences in manuscript submission behaviors, including risk aversion, perfectionism (Borghans et al., 2009; Closa et al., 2020; Djupe et al., 2019), and assessment of contribution (Lincoln et al., 2012; Wennerås & Wold, 2008). However, few large-scale studies have examined this question from the perspective of the authors. In this context, our paper seeks to address whether the sociodemographic characteristics of lead authors influence submission rates to elite journals. More specifically, based on a survey of more than 4,700 active authors (29.9 % women), we assess differences in gender, academic rank, and discipline in the likelihood of submitting to *Science, Nature*, and *PNAS* and the self-reported reasons for not submitting to these journals.

## Results

### Journal submission behavior

Women were less likely than men to report that they submitted a paper to *Science, Nature*, or *PNAS* (48.7% of men and 37.0% of women, see SI Table 7). These differences are statistically significant (see SI Table 8), with men, in general, being more likely to submit to *Science, Nature*, or *PNAS* than women while controlling for academic rank (OR=0.61, 95% CI[0.54, 0.70]). This relationship remains significant for each discipline (see Figure 1), with relatively higher percentages of men in each of the disciplines submitting to these journals (ranging from 44.2% to 53.4%), compared to women (ranging from 33.5% to 37.5%). We also investigated the mean number of manuscripts that respondents reported submitting to *Science, Nature*, or *PNAS* (see Figure 1 and SI Table 9). All disciplines combined, 552 women and 1,583 men submitted to these three elite journals. The mean was calculated by investigating only respondents who submitted at least one manuscript to *Science, Nature*, or *PNAS*. The number of submissions an author could have submitted depends on the discipline (due to the difference in publication behavior between disciplines) and career status (junior, senior, non-academic). In medical sciences (coefficient=-1.46, 95% CI[-2.14, -0.79], p=0.00) and natural sciences and engineering (coefficient=-0.80, 95% CI[-1.39, -0.22], p=0.01), women submitted fewer manuscripts than men. No statistically significant difference was observed for the social sciences (coefficient=-1.35, 95% CI[-3.00, 0.31], p=0.11, see SI Table 10).

**Figure 1.**
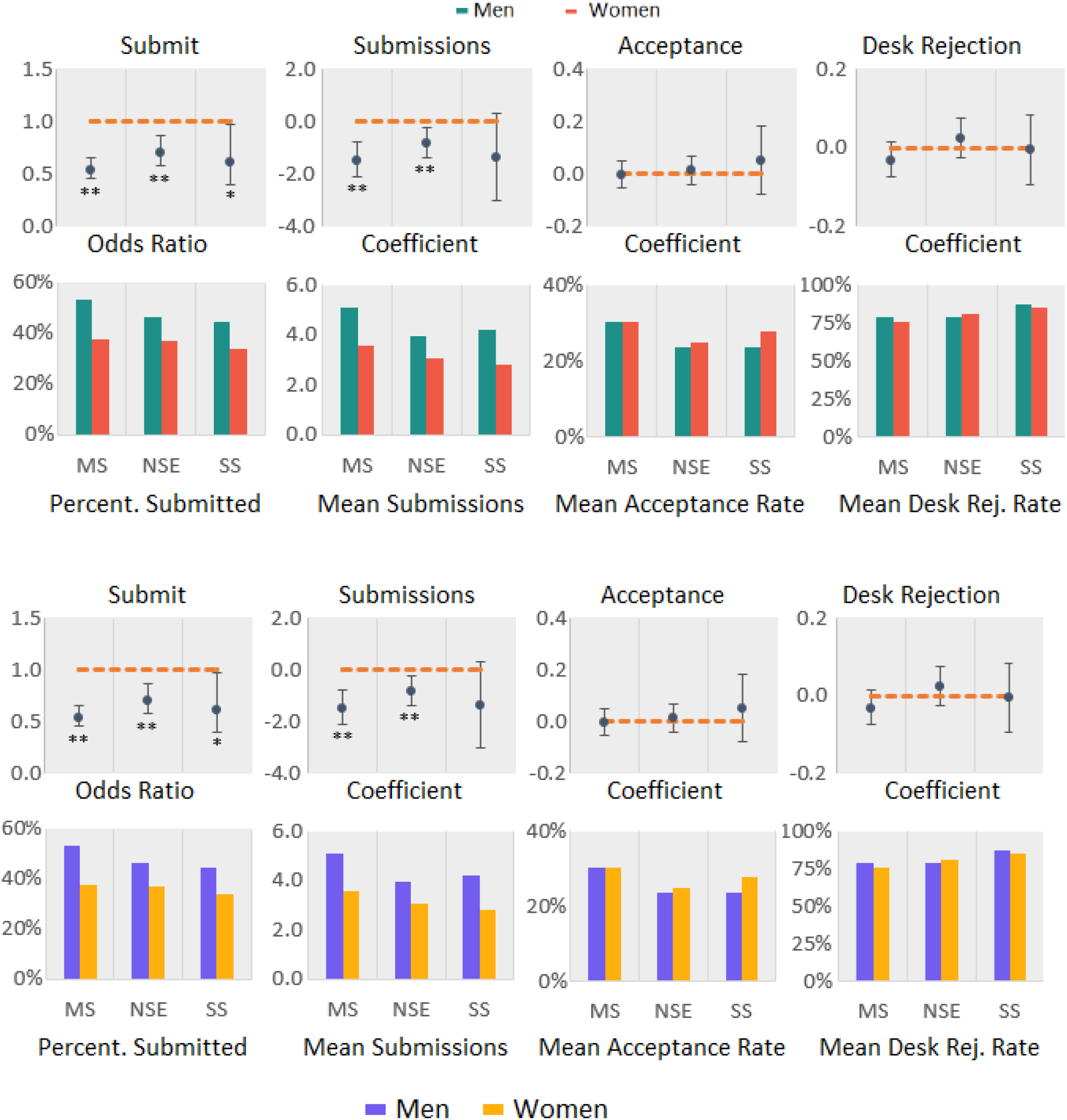
**Measures of submission rate, acceptance rate, and rejection rate with tests of statistical significance. Logistic regression was used for whether they submitted to an elit journal by individual discipline. Linear regression (mixed-effect model) was used understand the gender difference in number of submissions, acceptance rate and desk rejection rate to these journals by individual discipline. We controlled for rank in all regression analysis. Error bars represent the 95% confidence interval. All odds ratio and coefficient values are based on women over men. MS: Medical Sciences; NSE: Natural Sciences and Engineering; S.S.: Social Sciences. ** indicates p<0.01, * indicates p<0.05**

### Self-reported acceptance and rejection rate

We examined the difference between women and men regarding the self-reported acceptance and rejection rates for *Science, Nature*, and *PNAS*, using linear regression analysis while controlling for academic rank. The result shows no significant difference between women and men in getting their papers accepted (see Figure 1, SI Table 11, and SI Table 12). We also used a mixed-effect model to analyze the role of gender in the chances of getting manuscripts accepted, which yields similar conclusions to those above. We defined the desk rejection rate as the number of manuscripts that did not go out for peer review divided by the number of manuscripts submitted for each survey respondent. The relationship between desk rejection rate and gender were examined using linear regression and a mixed-effect model, again controlling for rank. Both models yield similar results: gender did not significantly impact the possibility of getting a desk rejection for manuscripts submitted to these journals, as illustrated in Figure 1 (as well as SI Table 13 and SI Table 14).

### Submission and desk rejections of authors highest cited papers

To understand whether the gender differences observed above also hold for the subset of higher impact research, we investigated respondents’ submission pattern for their most cited papers (not published in *Science, Nature, PNAS*, but also *Cell, Nature Communications, NEJM*, and *Science Advances*; see Methods and Materials for more details). Results show that a substantial proportion of the authors (48.6%) considered other journals than the one in which their paper was published, with no significant gender difference (see SI Table 17 and SI Table 18). However, only a small proportion of authors (9.1%) considered *Science, Nature*, or *PNAS*, again with no significant gender difference (see SI Table 19 and SI Table 20). As only a few authors considered publishing in these journals, it is not surprising to see that few authors submitted their most cited paper to *Science, Nature*, or *PNAS* (228 respondents), again with no significant gender gap (see SI Table 21 and SI Table 22). Similarly, no significant difference was observed in desk rejection rate for their most cited paper between men and women in any of the disciplines when submitting to *Science, Nature*, and *PNAS* (see SI Table 23 and SI Table 24). Submission behavior and the chance of getting a desk rejection for higher quality papers thus do not seem to differ significantly between men and women (Figure 2), although these conclusions are based on a very limited number of responses: 480 women and 1,404 men across all disciplines.

**Figure 2.**
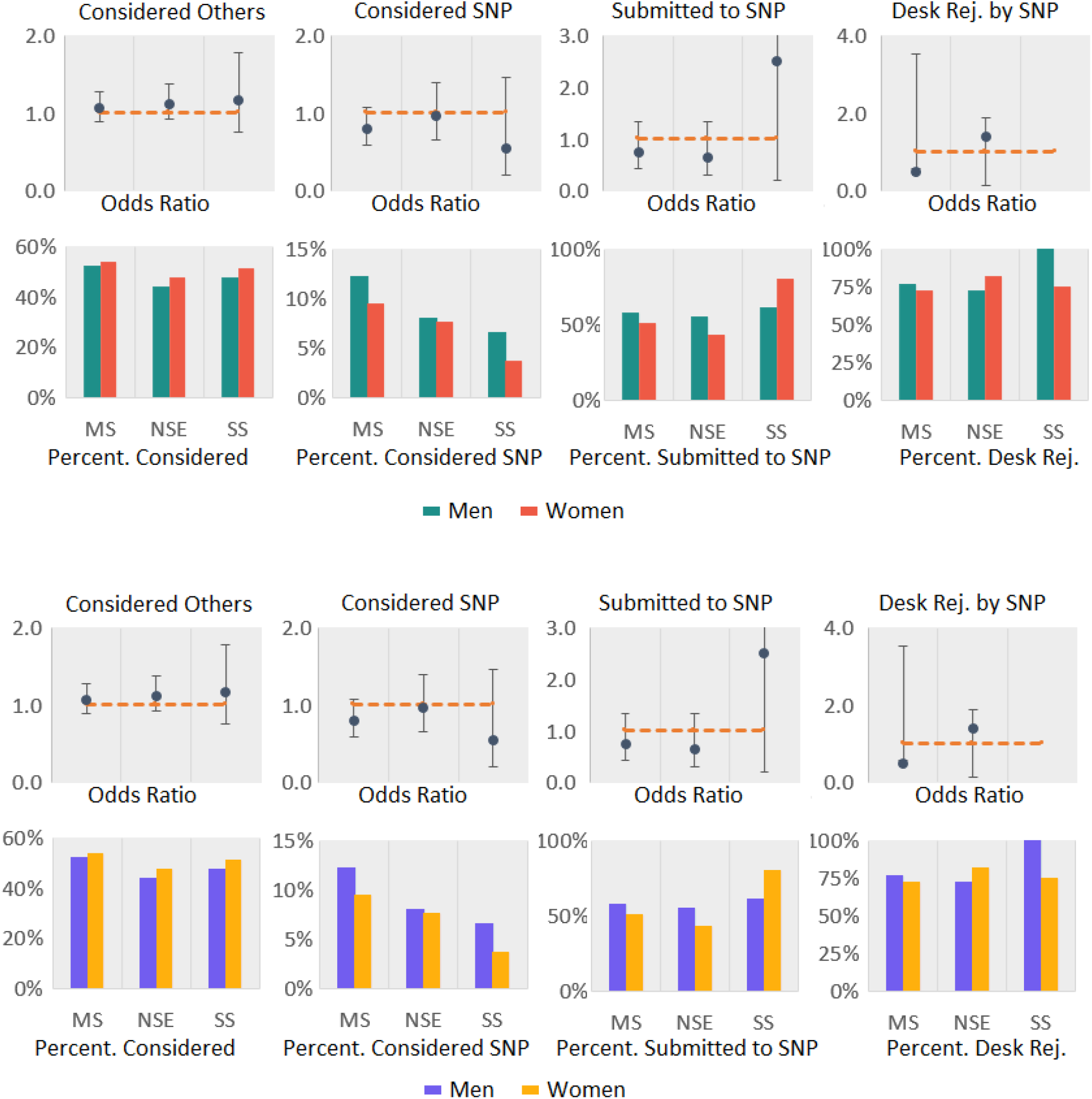
**Measures of journal consideration, submission rate, and desk rejection rate for most cited papers with odds ratios (women/men). Error bars represent the 95% confidence interval. We used logistic regression to analyze the relationship between gender and the four variables, while controlling for rank. Regression was done by each discipline separately. MS: Medical Sciences; NSE: Natural Sciences and Engineering; S.S.: Social Sciences. SNP: Science, Nature, PNAS.**

### Perception of the quality of research

Respondents were asked to rate their perception of the quality of their research (not their papers) in comparison to their peers on a five-point Likert scale ranging from Excellent (5) to Poor (1). Most respondents rated their work as good or excellent (86.7%), regardless of discipline, with very few rating their work as poor or fair (1.0%), and only a few respondents rated it as average (12.4%) (Figure 3 and SI Table 27). When considering responses regardless of discipline, no statistically significant difference was observed between men and women regarding how they rated the quality of their work (see SI Table 28). When disaggregating the respondents by discipline, it was found that compared with men, women in the natural sciences and engineering are more likely to rank their research quality lower (OR=0.83, 95% CI[0.67, 0.99], p=0.04). No statistically significant difference was observed in the medical (OR=1.18, 95% CI[0.99, 1.40], p=0.07) and social sciences (OR=1.13, 95% CI[0.93, 1.22], p=0.58)

**Figure 3.**
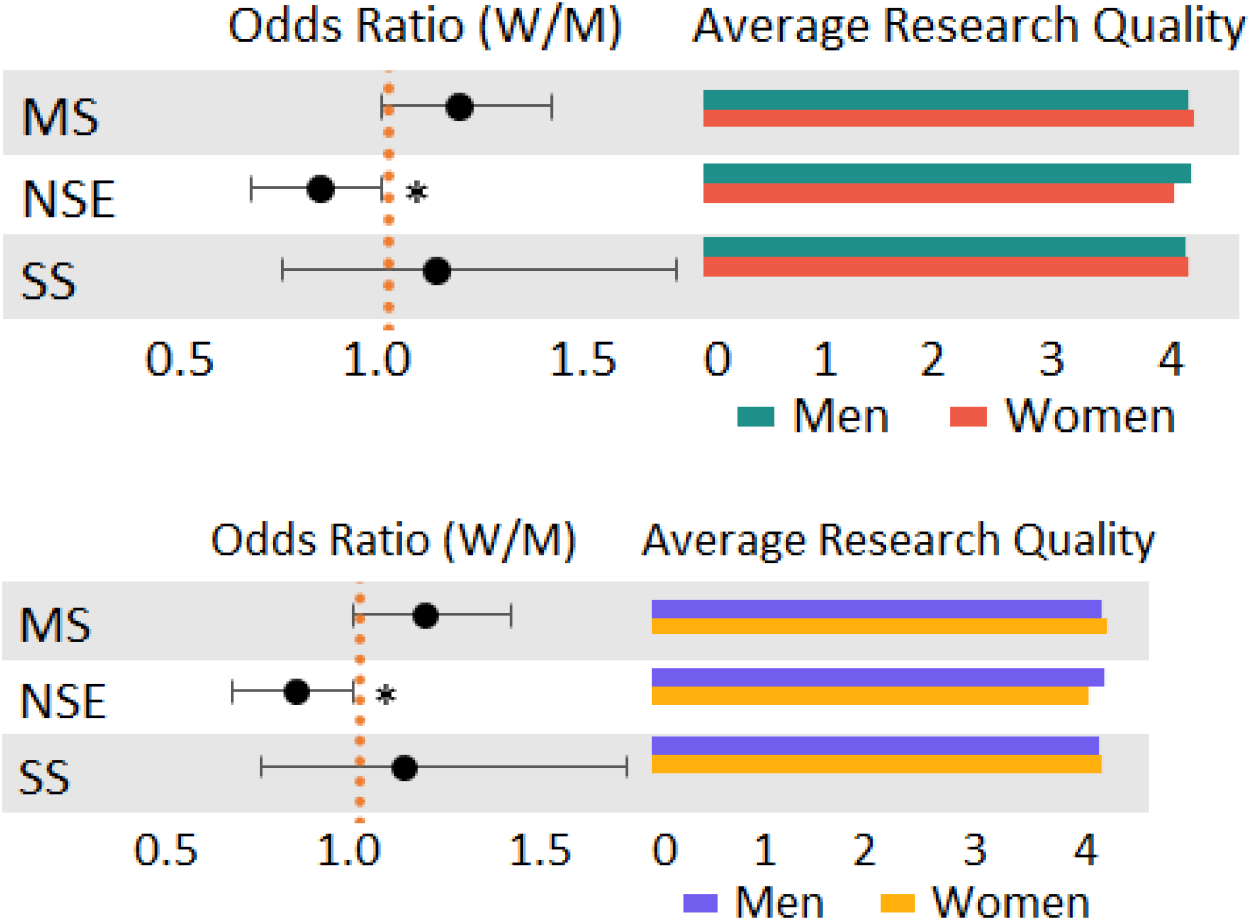
**Odds ratio (women/men) and mean value for research quality comparison with peers. Ordinal logistic regression was used to analyze the relationship between gender and the rating of research quality while controlling for respondents’ rank. Regression was done at the discipline level. Error bars represent the 95% confidence interval. MS: Medical Sciences; NSE: Natural Sciences and Engineering; S.S.: Social Sciences. *Indicates p<0.05**

### Reasons for not submitting to elite journals

We also investigated the authors’ rationale for why they did not submit their manuscripts (in general and their most cited) to the selected journals by asking respondents to select from a list of potential reasons. The results of the regression analysis are represented in Figure 4. None of the respondents with a known academic rank indicated that they did not submit to the journal because they were unaware of the journals. The most common reason men and women gave wa that their work would fit better in a more specialized journal.

**Figure 4.**
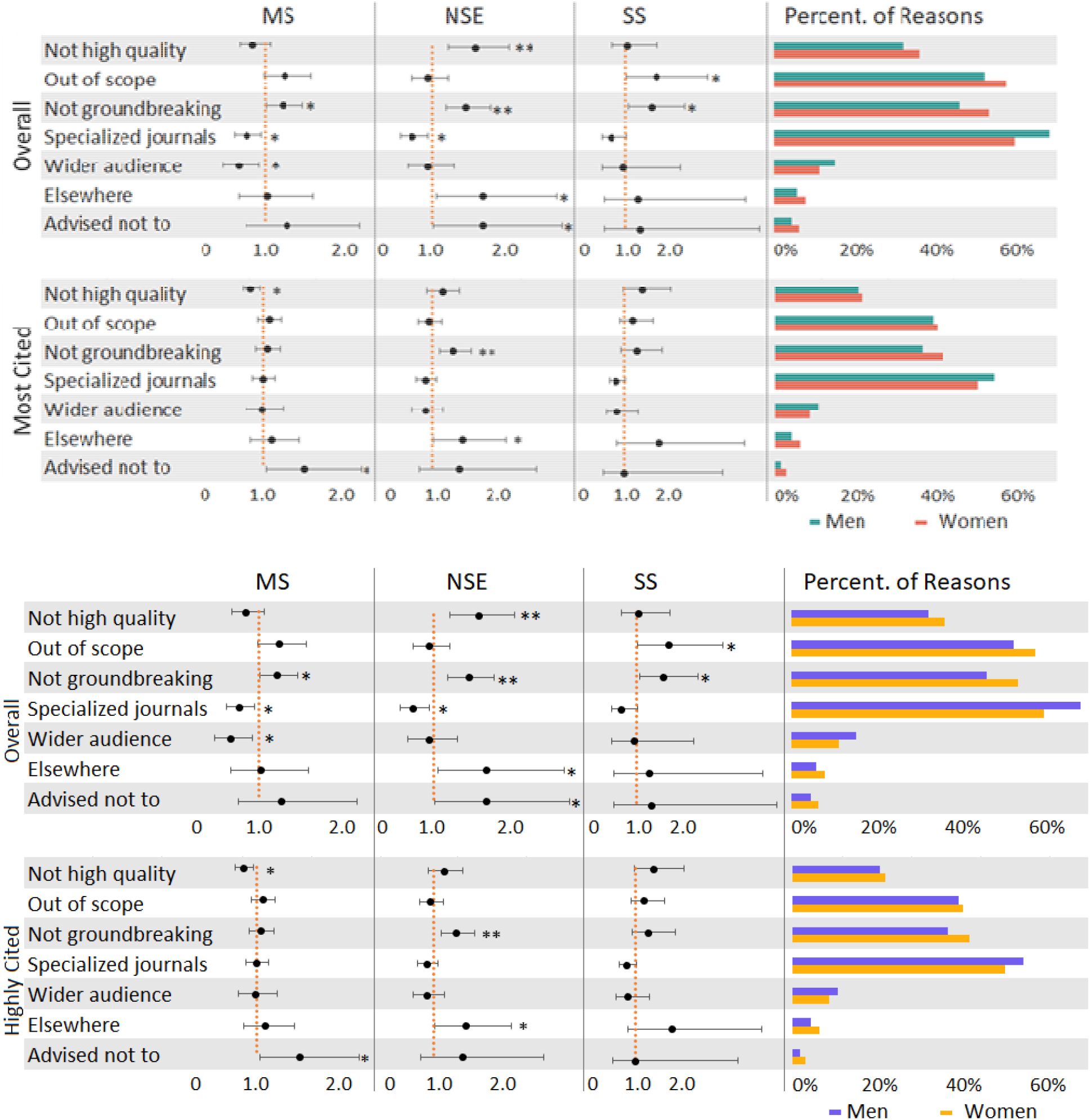
**Reasons for not submitting to top journals, overall and most cited paper, with odds ratios (women/men). Logistic regression was employed to analyze the relationship between gender and each of the reasons for not submitting. Regression was done by reason and discipline while controlling for rank. Error bars represent the 95% confidence interval. MS: Medical Sciences; NSE: Natural Sciences and Engineering; S.S.: Social Sciences. ** indicates p<0.01, * indicates p<0.05**

Two of the listed reasons are related to the quality and the novelty of the research. Women in natural sciences and engineering were more likely than men to indicate that “*the work was not of high enough quality*” as one of the reasons why they did not submit to *Science, Nature*, or *PNAS* (OR=1.59, 95% CI[1.22, 2.08], p=0.001). A similar finding was also found for the most cited paper, although only significant in the medical sciences (OR=0.75, 95% CI[0.60, 0.94], p=0.011). Regardless of discipline, women were more likely than men to (OR=1.26, 95% CI [1.07, 1.50]) indicate that they consider their “*work was not ground-breaking or sufficiently novel”* for the journal in question. For their most cited papers, it was also the case for authors publishing in natural sciences and engineering (OR=1.36, 95% CI[1.10, 1.67], p=0.004), as well as all disciplines combined (OR=1.19, 95% CI[1.04, 1.37], p=0.011) (see SI Table 16 and SI Table 26).

Reasons related to the scope and audience of the journals mostly show non-significant gender differences, with one exception. In general, men were more likely than women to indicate that the “*work would fit better in a more specialized journal”* for their general submissions (OR=0.72, 95% CI[0.60, 0.85], p=0.000). When disaggregated by discipline, a significant difference was observed both for the medical sciences (OR=0.74, 95% CI[0.57, 0.95], p=0.018) and for the natural sciences and engineering (OR=0.72, 95% CI[0.55, 0.95], p=0.018). However, no statistically significant difference was observed for the most cited papers. Men in the medical sciences were also more likely than women to indicate that the “*work would reach a wider audience in another journa*l” (OR=0.61, 95% CI[0.41, 0.92], p=0.017). No statistically significant difference was observed for the most cited papers. In contrast to the above results, women submitting in the social sciences were more likely to indicate that the “*work fell out of the scope of the journal*” (OR=1.84, 95% CI[1.03, 3.28], p=0.04).

The last two reasons are much less frequent and relate to the wishes of the co-authors and the advice received. In the natural sciences and engineering, women were more likely than men to indicate that their co-authors wished to submit elsewhere (OR=1.70, 95% CI[1.06, 2.74], p=0.029). This was also the case for their most cited paper (OR=1.57, 95% CI[1.02, 2.40], p=0.039). Without disaggregating by discipline, women were more likely to indicate that they did not submit their papers (in general and their most cited papers) to *Science, Nature*, or *PNAS* because they were advised not to. When disaggregating by discipline, this difference was also observed for respondents in natural sciences and engineering for their papers in general (OR=1.69, 95% CI[1.02, 2.81], p=0.042) and for medical sciences respondents for their most cited papers (OR=1.75, 95% CI[1.05, 2.93], p=0.032).

## Discussion and Conclusion

Women are particularly underrepresented in journals with the highest impact factor (Huang et al., 2020), with substantial consequences for their careers. While a large body of research has focused on the outcome and the process of peer review, fewer articles have explicitly focused on gendered submission behavior and the explanations for these differences. In our study of nearly five thousand active authors, we find that women are less likely to report having submitted papers and, when they have, to submit fewer manuscripts, on average, than men. This reinforces much of the literature demonstrating lower submission rates among women (Bendels et al., 2018; Grossman, 2020; Martinsen et al., 2022; Squazzoni et al., 2021). One strategy might be to optimize around homophily: women editors tend to disproportionately select women reviewers (Helmer et al., 2017), and women reviewers are more favorable to women-authored work (Murray et al., 2018). The positive reinforcement in the process will increase women reviewers and authors, which may influence submission behavior on the broader network. Another strategy might be to introduce quotas into peer review; however, this has been shown to have negative effects on women (Leibbrandt et al., 2018).

There is a demonstrated relationship between self-efficacy and publication output (Hemmings & Kay, 2009); however, our results on this were mixed. In the aggregate, no statistically significant difference was observed between men and women in how they rated the quality of their work. A difference was only observed in the natural sciences and engineering, where women ranked their research quality lower. The latter is particularly interesting, given that women tend to outperform in Engineering on several indicators (Ghiasi et al., 2015). The weak signal observed suggests that no particular intervention may be necessary to improve self-efficacy for this population; however, there may be some internalization of value premised on past experiences in peer review. Therefore, addressing larger issues of bias in evaluation (Lee et al., 2013) may serve to mitigate any gendered differences in self-efficacy.

Regardless of discipline, women were more likely than men to indicate that their “*work was not ground-breaking or sufficiently novel”* for the listed prestigious journals. Men were more likely than women to indicate that the “*work would fit better in a more specialized journal*.*”* Women were more likely to indicate that they did not submit their papers (in general and their most cited papers) to *Science, Nature*, or *PNAS* because they were advised not to. These results may reinforce the notion of risk aversion and perfectionism (Closa et al., 2020), suggesting that women have a higher internal standard for what constitutes novel research. However, the more alarming result is that women seem to be deterred from submitting from people within their scholarly network. One interpretation could be that the work is truly not sufficiently novel; this would suggest that men are innately predisposed to higher novelty research designs or that they are trained in this manner. Given no evidence of the former and sufficient evidence of differential mentoring for women (Nolan et al., 2008), it would stand to reason that greater attention should be made to equity in research training. However, the alternative interpretation is that women are not receiving the simple encouragement necessary to seek out higher-impact venues. Research administrators may seek simple interventions to increase this support. Journal editors may also want to consider dedicated outreach to women-led labs.

Addressing gender disparities in science requires a multifaceted approach that considers all components of the scientific system. In that context, we cannot overemphasize the need for a research evaluation reform—inclusing initiatives that span from the Leiden Manifesto, to the San Francisco Declaration on Research Assessment. But until those reforms are actually adopted by universities and funders and actions move beyond pledges, publishing remains a cornerstone of this system and central to research evaluation. Therefore, addressing mechanisms that create disparities in journal submission—be they self-imposed or otherwise—is essential for creating a robust and responsible research ecosystem.

## Materials and methods

A survey was conducted to explore the relationship between author gender and manuscript submission, rejection, and acceptance rates for high-impact, multidisciplinary journals in physical sciences, life sciences, and engineering (see S.I. appendix for the questionnaire). This study focuses on the results for *Science, Nature*, and *PNAS*. The target population consisted of active authors who published articles from 2008 to 2017 in journals indexed by WoS in medical sciences, natural sciences and engineering, and social sciences (Table S2). The questionnaire asked respondents to report on their manuscript submission experiences. It included a subsection focusing on their most cited paper published after 2010 not published in selected elite journals, i.e., *Science, Nature, PNAS* (which are analyzed in the main manuscript), as well as *Cell, Nature Communications, NEJM*, and *Science Advances* (which are analyzed in the S.I. appendix). Specific questions addressed their experiences submitting their most cited manuscripts to these journals, as well as their experiences submitting to these journals in general. Stata software (Standard Edition 17) was used to conduct the analyses, which consisted of applying ordinal logistic regression, multinomial logistic regression, and multiple logistic regression with a significance level or alpha of 0.05. Although we collected data on genders other than men and women, the independent variable studied is a binary gender variable (i.e., women or men), given the low number of respondents outside these two categories. Respondents were disaggregated by academic rank (i.e., junior, senior, or non-academic, excluding all unknowns) and discipline (i.e., social sciences, medical sciences, and natural sciences and engineering). More details about the regression analysis can be found in the appendix. The sample consisted of 6,002 respondents who participated in the survey, of whom 4,857 finished the questionnaire. An analysis of the attrition failed to identify a common point of departure, suggesting individual variability in dropout rather than failed survey construction. The final number of respondents in this study is 4,805, after the removal of 19 responses in Arts and Humanities and additional responses due to a lack of information for critical variables. Of the final number of respondents, 4,740 (98.6%) have a known academic rank and are included in the regression analysis (more details available in the S.I. appendix SI Table 6).

## Data availability

All data needed to evaluate the conclusions in the paper are present in the paper and the Supporting information. Aggregated,de-identified data by gender, discipline, and rank for analyses are available on GitHub (https://github.com/UWMadisonMetaScience/genderPublishing).

Due to Institutional Review Board restrictions and the terms of participant consent, the survey data used in this study cannot be shared publicly. However, to support transparency and reproducibility, all analysis code and a synthetic dataset—constructed with randomly generated values but matching the original data structure—are available on Zenodo: https://doi.org/10.5281/zenodo.16327580.

## Acknowledgments

Vincent Larivière acknowledges funding from the Canada Research Chairs program. The authors would like to thank the survey respondents and the Indiana University Center for Survey Research for their critical contribution to the study. This analysis was approved by McGill University’s Research Ethics Board II, R.E.B. File #: 501-0518 *Factors affecting scientific manuscript submission and acceptance rates*.

## Notes

### Competing Interest Statement

The authors have declared no competing interest.

### Summary of Updates

Minor edits related to colors in figures and conclusions.

