## Supplementay materials for "Gender differences in submission behavior exacerbate publication disparities in elite journals"

### SUPPORTING INFORMATION

#### Expansion on Materials and Methods

##### Sampling frame construction

The sampling frame was constructed using data extracted from Web of Science (WoS), as hosted by the Centre for Science and Technology Studies (CWTS) at Leiden University. Using the author disambiguation algorithm developed by Caron and van Eck (2014)[1], we extracted metadata of all publications for authors who have been corresponding authors of at least one paper between 2008 and 2017. The metadata included the email addresses of these authors; these email addresses were used to contact the participant population. A minimum cut-off point of 10 papers for the identification of prolific authors was selected. When considering whether an author is a prolific author, only research papers were considered. The total study population consists of 494,781 prolific authors, data on the gender and discipline in which they publish were known for 415,697 authors (84.0%). The gender of an author for the sampling frame was determined by the algorithm developed in Larivière et al. (2013)[2] whereas the gender variable used in the analysis were determined by the responses of the authors. The discipline in which an author publishes was determined by the WoS journal classification.

##### Identifying most cited papers

Several questions in the questionnaire asks the authors about their decisions pertaining to their most cited paper. The questionnaires are customized to each respondent, showing them the title of their most cited paper when these questions are asked. The most cited paper was identified from the previous corpus of papers by the author published between 2008 and 2017, that has received the most citations at the time of data extraction and were not published in the selected elite journals. Despite the fact that citations take time to accumulate, the skewed nature of citation distributions—where most papers remain lowly cited, and a few papers get highly cited—leads to a situation where most cited papers are not necessarily the oldest papers published by respondents.

##### Elite journal selection

The questionnaire mainly consists of questions related to the author's submission behavior as they pertain to elite journals. The seven journals were chosen given their general reputation and research impact. All seven journals were in the top five journals with the highest impact factors in the 2016

[1] Caron, E. & Van Eck, N.J. (2014). "Large scale author name disambiguation using rule-based scoring and clustering." in *Proceedings of the 19th international conference on science and technology indicators*, Leiden.

[2] Larivière, V., Ni, C., Gingras, Y., Cronin, B. & Sugimoto, C.R. (2013). "Bibliometrics: Global gender disparities in science." *Nature*, 504: 211–213. doi:10.1038/504211a.

Journal Citation Reports categories of "Multidisciplinary sciences", "General & internal medicine," and "Biology." Journal Citation Reports[3] (https://jcr.clarivate.com/jcr/) is a resource published annually by Clarivate Analytics that provides the impact factor for all journals included in WoS. The journal impact factor[4] is a calculation that employs citations that articles in a journal received as an indication of the influence of that journal in its field.

#### Regression Analysis

This study used regression analysis to analyze the data. The analysis described here is similar to Ni et. al., (2021), [5] and how it is reported in its supplementary materials. Here the discussion is slightly adapted to reflect the variables and questions used within this study. The regression procedures include binary logistic regression, ordinal logistic regression, linear regression and mix-effect model. Regression analysis is usually used to explore relationships between dependent and independent variables. The most common is linear regression, in which:

$$Y_i = \beta_0 + \beta_1 X_i + \beta_2 Z_i + \ldots + e_i$$

where with one unit change in X there is $\beta_1$ differences in Y after Z and other variables are controlled. However, this approach assumes that all dependent variables are normally distributed. Furthermore, it often assumes that dependent variable Y is continuous. Our dataset, like many social science surveys, is replete with categorical variables. We want to understand whether there is a gendered difference when selecting one category over another. Therefore, we turned to logistic regression. Specific procedures and analysis methods vary by the scale of dependent variables, as well as the number of variable categories.

For questions with answers measured in ordinal scales (e.g., Likert Scaling), ordinal logistic regression analysis was employed. For questions with answers (the dependent variables) presented in a categorical, unordered scale, binary logistic regression (when the dependent variable only contains two categories) was performed. The point statistic was still the odds ratio of women over men. Here again, gender is the independent variable, with the academic rank being the controlled variable in the case when a dependent variable only has two categories (e.g., "yes" or "no" to an answer), binary logistic regression analysis was conducted to analyze the data. An odds ratio value significantly over 1 indicates that women were more likely than men to select "Yes" for this question. A separate regression analysis is performed for each discipline with rank being controlled. For analyses concerning all disciplines, we control for both academic rank and discipline.

---

[3] More information at https://clarivate.libguides.com/jcr.

[4] Garfield, E. (2006). "The history and meaning of the journal impact factor." *JAMA*, 295(1):90–93. doi:10.1001/jama.295.1.90

[5] Ni, C., Smith, E., Yuan, H., Larivière, V., & Sugimoto, C.R. (2021). "The gendered nature of authorship." *Science Advances*, 7(36). doi:10.1126/sciadv.abe4639.

The binary logistic regression model for each discipline can be described as follows:

$Y_i = \text{Log}\left(\frac{p}{1-p}\right) = \beta_0 + \beta_1 X_i + \beta_2 Z_i + \ldots + e_i$

where $Y_i$ is the log probability of the binary dependent variable (such as whether you submitted to a journal), $X_i$ is the binary independent variable gender (woman=1 and man=0), and $Z_i$ is the other control variable(s).

The mixed-effect linear model for each discipline can be described as follows:

$$Y_i = \beta_0 + \beta_1 X_i + Z_i + \ldots + e_i$$

Where $Y_i$ is the dependent variable (number of submissions, acceptance rate, or rejection rate), $X_i$ is the gender of the individual (the fixed effect), and $Z_i$ is the rank of the individual (random effect variable).

Recoded categories

The academic rank variable was created by consulting the respondents' answers on the questions regarding academic rank (for those respondents that indicated their primary sector affiliation as Academia), and position (for those respondents from the Government, Privaye or Not_for_Profit sector). Responses were recategorized into junior, senior,and non-academic, with unknowns removed from analyses, as per SI Table 1:

**SI Table 1 Recategorized academic rank.**

| Questionnaire categories | New categories | Count |
|---|---|---|
| Graduate student | junior | 991 |
| Postdoctoral fellow | | |
| Research Associate | | |
| Assistant Professor | | |
| Associate Professor | senior | 3,012 |
| Full Professor | | |
| Emeritus Professor | | |
| non-academic respondents | non-academic | 737 |
| No answer and other | unknowns | 65 |

Due to limited responses, analysis per journal was not always viable. The results pertaining to the journals were aggregated, with new categories based on the shared similarities in disciplinary foci of the journals and their prestige. The categories are as follow:

- Science, Nature, and PNAS (S.N.P.)
- Nature Communications and Science Advances (NC.SA.)

- NEJM and Cell (NEJM.C.)

Similarly, the various options for research areas in the questionnaire were reclassified into three broader areas that share common research and dissemination practices, namely natural science and engineering (NSE), medical science (MS), and social sciences (SS), as shown in SI Table 2.

**SI Table 2 Recategorized research areas to disciplines.**

| Disciplinary area | Field | Surveyed | Respondents | Analytical Sample |
|---|---|---|---|---|
| Medical Sciences | Biomedical Research | 63,335 | 832 | 678 |
| | Clinical Medicine | 152,056 | 1,394 | 1,134 |
| | Health | 12,807 | 245 | 208 |
| Natural Sciences and Engineering | Biology | 47,539 | 670 | 543 |
| | Chemistry | 36,830 | 395 | 321 |
| | Earth and Space | 36,573 | 576 | 467 |
| | Engineering and Technology | 61,746 | 559 | 406 |
| | Mathematics | 18,674 | 275 | 206 |
| | Physics | 46,694 | 592 | 475 |
| Social Sciences | Professional Fields | 3,480 | 82 | 66 |
| | Psychology | 9,885 | 243 | 207 |
| | Social Sciences | 4,354 | 114 | 94 |
| Arts and Humanities | Arts | 291 | 9 | / |
| | Humanities | 475 | 14 | / |
| Unknown | Unknown | 42 | 2 | 0 |
| | **Total** | **494,781** | **6,002** | **4,805** |

<u>Description of the analytical sample</u>

The Indiana University Centre for Survey Research (CSR) was contracted to administer the survey. The description of the survey response is based on the technical report produced by the CSR. After data cleaning (e.g., removing duplicates), a total of 489,777 email invitations were sent, and 6,002 individuals participated in the online survey. The responses consisted of 4,857 completed questionnaires. After reviewing the complete questionnaires, responses of authors in Arts (8) and humanities (11) questionnaires were excluded. Lastly, we further removed additional responses due to lack of information for critical variables (e.g. gender, and discipline). An analysis of the attrition failed to identify a common point of departure, suggesting individual variability in dropout rather than failed survey construction. The final analytical sample consists of 4,805 respondents. Depending on the variable, the number of responses for analyzing them may vary.

The response rate of the survey was 1.2%, as classified according to The American Association for Public Opinion Research. 2015. Standard Definitions: Final Dispositions of Case Codes and Outcome Rates for Surveys. 8th edition (AAPOR), as summarized in SI Table 3.

**SI Table 3 Summary of response rate of questionnaire (adapted from McGill University – Gender Ethnicity Publishing Study Methods Summary technical report[6]).**

| Disposition | Definition | Count | Response rate (RR2) |
|---|---|---|---|
| Complete (I) | Respondent completed survey | 4,944 | 1.2% |
| Partial (P) | Respondent answered at least one question item but did not complete survey. | 1,058 | |
| Implicit Refusal (R) | Respondent clicked survey link but did not answer any items | 1,655 | |
| Nothing Returned (UH) | Respondent did not respond to survey; it is unknown if any email messages were read. | 374,341 | |
| Undeliverable (UO) | Recruitment message was not received by intended recipient due to email and/or mailing returns | 108,163 | |
| Not Eligible | Sample member indicated they were not eligible to participate; also includes cases of authors who have deceased since their publication listed in the database | 58 | |
| Total | | 490,219 | |

The following tables (SI Table 4, SI Table 5, SI Table 6) provide a description of the population, respondents, and analytical sample by gender.

**SI Table 4 Gender composition of population, respondents, and analysis sample.**

| Domain | Population / Surveyed | | Respondents | | Analytical sample | |
|---|---|---|---|---|---|---|
| | N | % | N | % | N | % |
| Men | 286,571 | 57.9% | 3,508 | 58.4% | 2,832 | 58.9% |
| Women | 129,592 | 26.2% | 1,774 | 29.6% | 1,435 | 29.9% |

6 Indiana University Centre for Survey Research (2019). McGill University – Gender Ethnicity Publishing Study Methods Summary. Technical Report. Unpublished.

| | | | | | | |
|---|---|---|---|---|---|---|
| Unknown | 78,618 | 15.9% | 720 | 12.0% | 538 | 11.2% |
| All | 494,781 | 100.0% | 6,002 | 100.0% | 4,805 | 100.0% |

**SI Table 5 Discipline composition of population, respondents, and analysis sample.**

| Domain | Population / Surveyed | | Respondents | | Analytical sample | |
|---|---|---|---|---|---|---|
| | N | % | N | % | N | % |
| MS | 228,198 | 46.1% | 2,471 | 41.2% | 2,020 | 42.0% |
| SS | 17,719 | 3.6% | 439 | 7.3% | 366 | 7.6% |
| NSE | 248,098 | 50.1% | 3,069 | 51.1% | 2,419 | 50.3% |
| AH | 766 | 0.2% | 23 | 0.4% | | 0.0% |
| All disciplines | 494,781 | 100.0% | 6,002 | 100.0% | 4,805 | 100.0% |

**SI Table 6 Discipline, gender, and academic rank composition of analytical sample.**

| | MS (N=2,020) | | | | NSE (N=2,419) | | | | SS (N=366) | | | |
|---|---|---|---|---|---|---|---|---|---|---|---|---|
| Rank/Role | Women | | Men | | Women | | Men | | Women | | Men | |
| | N | % | N | % | N | % | N | % | N | % | N | % |
| Junior | 200 | 24.5% | 228 | 18.9% | 147 | 27.6% | 360 | 19.1% | 31 | 18.7% | 25 | 12.5% |
| Senior | 465 | 57.0% | 762 | 63.3% | 281 | 52.7% | 1,219 | 64.6% | 124 | 74.7% | 161 | 80.5% |
| Non-academic | 131 | 16.1% | 189 | 15.7% | 100 | 18.8% | 294 | 15.6% | 10 | 6.0% | 13 | 6.5% |
| Unknown | 20 | 2.5% | 25 | 2.1% | 5 | 0.9% | 13 | 0.7% | 1 | 0.6% | 1 | 0.5% |

#### Limitations

Several limitations should be noted regarding the findings of this study. Firstly, the survey response rate is less than 2%, which may limit the generalizability of the findings. In addition, asking researchers about their behaviour and intentions regarding their manuscript submissions retroactively can be complicated. There is a concern related to the capacity of researchers to remember their motivations at the time of submission; researchers with longer careers—and therefore a higher number of papers—may have submitted to various journals for various reasons. This would also make it difficult to ascertain the reasons why they do not submit to the journals, as the motivations would be different for each paper. Moreover, reasons for not submitting to these journals were not exhaustive, as noted by the various comments made by respondents. One of the notable reasons given by respondents that relate to the gender component of the study is that they did not submit to these journals due to the low percentage of women on the journal editorial boards. Based on respondents' comments, page fees and article processing charges (APC) were also a serious consideration for some authors when deciding where to publish their work. Unfortunately, this consideration was not incorporated into the listed reasons in the questionnaire.

There are also several factors pertaining to the nature of most cited papers that complicates the interpretation of the results. For example, some respondents stated that their most cited paper was an invited paper of some kind, thus the choice of journal was already made for them and, therefore, the survey questions were irrelevant. In addition, even though their most cited paper was used as a proxy of a high-quality paper, the paper in question was not necessarily a highly cited paper, or necessarily significantly more cited than their other papers, or a paper of high quality (e.g., all their other papers could have been cited once and their most cited paper twice). That being said, the papers analyzed had, on average, a research impact 2.3 times higher than the world average of their field, which shows that they can be categorized as high impact. On the whole, this was the best proxy available for assignment of high quality papers to researchers and the effect is reduced by only investigating the subset of prolific researchers.

Some limitations relate to the passage of time between manuscript submission and data collection. Academic rank refers to the academic rank of the author at the time of the survey and not at the time of manuscript submission. This could result in a scenario where a respondent's highest cited paper could have been submitted in 2008 while they were a junior academic but in 2019, at the time of the survey, they have since changed academic rank to senior academic. Therefore, this should be interpreted as the maximu rank reached by the respondent. Review policies of journals are also a potential limitation as data pertaining to the review policies at the time of manuscript submission were not collected. If a journal had a double-blind review policy at the time of submission, it might reduce potential gender bias and be a confounding effect if journals of different peer review types were grouped together. It is likely that the review policy of the journals were single-blind at the time of submission of the papers as it is the review policy most used in the fields of life sciences, physical sciences, and engineering[7] and thus unlikely to have changed. While this could be a factor when we investigated acceptance rate, it is not a consideration when investigating desk rejections as none of the journals include an editor blinded review policy.

[7] Ware, M. (2008). "Peer review in scholarly journals: Perspective of the scholarly community — an international study." *Information Services & Use*, 28(2). doi:10.5555/1454388.1454399.

Regression and additional descriptive tables

The questionnaire was divided into four parts and the following sections are ordered by the sections of the questionnaire. The sections are as follows:

1. PART 1: Consent for Research Study Participation (information regarding questionnaire – no questions)
2. PART 2: Your manuscript submissions (i.e., questions pertaining to submissions in general)
3. PART 3: Your journal article (i.e., questions pertaining to most cited paper)
4. PART 4: Demographics (as well as a question pertaining to quality of research in general and open comments)

PART 2: Submissions in general

Journal submission behavior

**SI Table 7 Question: "For each of the journals listed in the table below, please indicate the appropriate number of manuscripts submitted" – percentage by discipline, journal, and gender ever submitted.**

| S.N.P. | Women | | | | Men | | | |
|---|---|---|---|---|---|---|---|---|
| | No | Yes | No Response | %Yes | No | Yes | No Response | %Yes |
| MS | 504 | 303 | 9 | 37.5% | 555 | 636 | 13 | 53.4% |
| NSE | 328 | 194 | 11 | 37.2% | 1,000 | 860 | 26 | 46.2% |
| SS | 109 | 55 | 2 | 33.5% | 110 | 87 | 3 | 44.2% |
| All disciplines | 941 | 552 | 22 | 37.0% | 1,665 | 1 583 | 42 | 48.7% |
| | Women | | | | Men | | | |
| NC.SA. | No | Yes | No Response | %Yes | No | Yes | No Response | %Yes |
| MS | 670 | 137 | 9 | 17.0% | 902 | 283 | 19 | 23.9% |
| NSE | 429 | 91 | 13 | 17.5% | 1,476 | 375 | 35 | 20.3% |
| SS | 158 | 5 | 3 | 3.1% | 181 | 14 | 5 | 7.2% |
| All disciplines | 1,257 | 233 | 25 | 15.6% | 2,559 | 672 | 59 | 20.8% |
| | Women | | | | Men | | | |
| NEJM.C | No | Yes | No Response | %Yes | No | Yes | No Response | %Yes |
| MS | 603 | 204 | 9 | 25.3% | 795 | 390 | 19 | 32.9% |
| NSE | 505 | 12 | 16 | 2.3% | 1,816 | 34 | 36 | 1.8% |
| SS | 152 | 11 | 3 | 6.7% | 182 | 13 | 5 | 6.7% |
| All disciplines | 1,260 | 227 | 28 | 15.3% | 2,793 | 437 | 60 | 13.5% |

**SI Table 8 Question: "For each of the journals listed in the table below, please indicate the appropriate number of manuscripts submitted" – odds ratio (women to men) values for the probability of submitting.**

| Journal | Discipline | Odds ratio | Std. Err. | z | p>\|z\| | 95% CI Lower | 95% CI Upper | obs |
|---|---|---|---|---|---|---|---|---|
| S.N.P. | MS | 0.54 | 0.05 | -6.45 | 0.000 | 0.45 | 0.65 | 1,937 |
| | NSE | 0.70 | 0.07 | -3.40 | 0.001 | 0.58 | 0.86 | 2,356 |
| | SS | 0.62 | 0.14 | -2.13 | 0.033 | 0.40 | 0.96 | 355 |
| | All disciplines | 0.61 | 0.04 | -7.32 | 0.000 | 0.54 | 0.70 | 4,648 |
| NC.SA. | MS | 0.65 | 0.08 | -3.68 | 0.000 | 0.52 | 0.82 | 1,931 |
| | NSE | 0.81 | 0.11 | -1.60 | 0.110 | 0.63 | 1.05 | 2,346 |
| | SS | 0.35 | 0.19 | -1.91 | 0.056 | 0.12 | 1.02 | 352 |
| | All disciplines | 0.70 | 0.06 | -4.06 | 0.000 | 0.59 | 0.83 | 4,629 |
| NEJM. C. | MS | 0.71 | 0.07 | -3.35 | 0.001 | 0.58 | 0.87 | 1,931 |
| | NSE | 1.22 | 0.42 | 0.59 | 0.558 | 0.63 | 2.39 | 2,342 |
| | SS | 1.08 | 0.46 | 0.18 | 0.859 | 0.47 | 2.50 | 352 |

| | | | | | | | | |
|---|---|---|---|---|---|---|---|---|
| | All disciplines | 0.75 | 0.07 | -2.95 | 0.003 | 0.62 | 0.91 | 4,625 |

**SI Table 9 Question: "For each of the journals listed in the table below, please indicate the appropriate number of manuscripts submitted" – mean number of manuscript submissions**

| Average number of submissions to S.N.P. | Women | | | | | Men | | | | |
|---|---|---|---|---|---|---|---|---|---|---|
| | Obs | Mean | Std. dev. | Min | Max | Obs | Mean | Std. dev. | Min | Max |
| MS | 303 | 3.60 | 3.51 | 1 | 34 | 636 | 5.12 | 5.41 | 1 | 35 |
| NSE | 194 | 3.04 | 2.38 | 1 | 13 | 860 | 3.95 | 3.95 | 1 | 30 |
| SS | 55 | 2.80 | 3.36 | 1 | 19 | 87 | 4.22 | 5.49 | 1 | 30 |
| All disciplines | 552 | 3.32 | 3.15 | 1 | 34 | 1,583 | 4.43 | 4.71 | 1 | 35 |
| Average number of submissions to NC.SA. | Women | | | | | Men | | | | |
| | Obs | Mean | Std. dev. | Min | Max | Obs | Mean | Std. dev. | Min | Max |
| MS | 137 | 1.74 | 1.34 | 1 | 10 | 281 | 1.90 | 1.24 | 1 | 8 |
| NSE | 89 | 1.65 | 1.08 | 1 | 6 | 374 | 1.86 | 1.57 | 1 | 17 |
| SS | 5 | 1.00 | 0.00 | 1 | 1 | 14 | 1.71 | 1.14 | 1 | 5 |
| All disciplines | 231 | 1.69 | 1.23 | 1 | 10 | 14 | 1.71 | 1.14 | 1 | 5 |
| Average number of submissions to NEJM.C. | Women | | | | | Men | | | | |
| | Obs | Mean | Std. dev. | Min | Max | Obs | Mean | Std. dev. | Min | Max |
| MS | 203 | 1.89 | 1.34 | 1 | 10 | 389 | 2.25 | 2.00 | 1 | 16 |
| NSE | 12 | 1.67 | 1.44 | 1 | 6 | 34 | 1.76 | 1.48 | 1 | 8 |
| SS | 11 | 1.64 | 0.92 | 1 | 4 | 13 | 1.15 | 0.38 | 1 | 2 |
| All disciplines | 226 | 1.86 | 1.32 | 1 | 10 | 436 | 2.18 | 1.95 | 1 | 16 |

**SI Table 10 Question: "For each of the journals listed in the table below, please indicate the appropriate number of manuscripts submitted" – mixed linear regression.**

| Journal | Discipline | Coefficient | Std. dev. | t | p>\|t\| | 95% CI Lower | 95% CI Upper | obs |
|---|---|---|---|---|---|---|---|---|
| S.N.P. | MS | -1.46 | 0.34 | -4.26 | 0.000 | -2.14 | -0.79 | 921 |
| | NSE | -0.80 | 0.30 | -2.69 | 0.007 | -1.39 | -0.22 | 1,042 |

| | | | | | | | | |
|---|---|---|---|---|---|---|---|---|
| | SS | -1.35 | 0.84 | -1.61 | 0.110 | -3.00 | 0.31 | 140 |
| | All disciplines | -1.19 | 0.22 | -5.41 | 0.000 | -1.62 | -0.76 | 2,103 |
| NC.SA. | MS | -0.19 | 0.13 | -1.41 | 0.161 | -0.45 | 0.08 | 411 |
| | NSE | -0.19 | 0.18 | -1.09 | 0.276 | -0.54 | 0.16 | 459 |
| | SS | -0.71 | 0.54 | -1.32 | 0.207 | -1.86 | 0.44 | 19 |
| | All disciplines | -0.20 | 0.11 | -1.89 | 0.059 | -0.42 | 0.01 | 889 |
| NEJM.C. | MS | -0.36 | 0.15 | -2.39 | 0.017 | -0.65 | -0.06 | 575 |
| | NSE | -0.08 | 0.50 | -0.17 | 0.867 | -1.10 | 0.93 | 46 |
| | SS | 0.59 | 0.32 | 1.84 | 0.081 | -0.08 | 1.26 | 24 |
| | All disciplines | -0.31 | 0.14 | -2.23 | 0.026 | -0.58 | -0.04 | 645 |

<u>Acceptance rate</u>

**SI Table 11 Question: "For each of the journals listed in the table below, please indicate the appropriate number of manuscripts accepted" – mean acceptance rate**

| | Women | | | | | Men | | | | |
|---|---|---|---|---|---|---|---|---|---|---|
| Acceptance Rate of S.N.P. | Obs | Mean | Std. dev. | Min | Max | Obs | Mean | Std. dev. | Min | Max |
| MS | 262 | 0.30 | 0.35 | 0 | 1 | 565 | 0.30 | 0.34 | 0 | 1 |
| NSE | 177 | 0.25 | 0.35 | 0 | 1 | 798 | 0.24 | 0.34 | 0 | 1 |
| SS | 53 | 0.28 | 0.38 | 0 | 1 | 80 | 0.24 | 0.36 | 0 | 1 |

| | | | | | | | | | | |
|---|---|---|---|---|---|---|---|---|---|---|
| All disciplines | 492 | 0.28 | 0.35 | 0 | 1 | 1,443 | 0.26 | 0.34 | 0 | 1 |
| Acceptance Rate of NC.SA. | Women | | | | | Men | | | | |
| | Obs | Mean | Std. dev. | Min | Max | Obs | Mean | Std. dev. | Min | Max |
| MS | 137 | 0.27 | 0.39 | 0 | 1 | 281 | 0.32 | 0.39 | 0 | 1 |
| NSE | 89 | 0.30 | 0.41 | 0 | 1 | 374 | 0.36 | 0.42 | 0 | 1 |
| SS | 5 | 0.20 | 0.45 | 0 | 1 | 14 | 0.24 | 0.42 | 0 | 1 |
| All disciplines | 231 | 0.28 | 0.40 | 0 | 1 | 669 | 0.34 | 0.41 | 0 | 1 |
| Acceptance Rate of NEJM.C. | Women | | | | | Men | | | | |
| | Obs | Mean | Std. dev. | Min | Max | Obs | Mean | Std. dev. | Min | Max |
| MS | 202 | 0.27 | 0.38 | 0 | 1 | 388 | 0.26 | 0.36 | 0 | 1 |
| NSE | 12 | 0.17 | 0.33 | 0 | 1 | 34 | 0.18 | 0.34 | 0 | 1 |
| SS | 11 | 0.39 | 0.47 | 0 | 1 | 13 | 0.04 | 0.14 | 0 | 0.5 |
| All disciplines | 225 | 0.27 | 0.38 | 0 | 1 | 435 | 0.25 | 0.36 | 0 | 1 |

**SI Table 12 Question: "For each of the journals listed in the table below, please indicate the appropriate number of manuscripts accepted" – linear regression**

| Journal | Discipline | Coefficient | Std. dev. | t | p>\|t\| | 95% CI Lower | 95% CI Upper | obs |
|---|---|---|---|---|---|---|---|---|
| S.N.P. | MS | 0.00 | 0.03 | 0.01 | 0.989 | -0.05 | 0.05 | 810 |
| | NSE | 0.01 | 0.03 | 0.48 | 0.628 | -0.04 | 0.07 | 964 |
| | SS | 0.05 | 0.07 | 0.78 | 0.435 | -0.08 | 0.18 | 132 |
| | All disciplines | 0.01 | 0.02 | 0.57 | 0.566 | -0.03 | 0.05 | 1,906 |
| NC.SA. | MS | -0.05 | 0.04 | -1.15 | 0.252 | -0.13 | 0.03 | 411 |

| | | | | | | | | |
|---|---|---|---|---|---|---|---|---|
| | NSE | -0.07 | 0.05 | -1.42 | 0.156 | -0.17 | 0.03 | 459 |
| | SS | -0.04 | 0.22 | -0.16 | 0.877 | -0.51 | 0.44 | 19 |
| | All disciplines | 0.06 | 0.03 | -1.78 | 0.075 | -0.12 | 0.01 | 889 |
| NEJM.C. | MS | 0.02 | 0.03 | 0.65 | 0.517 | -0.04 | 0.08 | 573 |
| | NSE | -0.02 | 0.11 | -0.15 | 0.885 | -0.25 | 0.21 | 46 |
| | SS | 0.43 | 0.15 | 2.79 | 0.011 | 0.11 | 0.74 | 24 |
| | All disciplines | 0.03 | 0.03 | 1.07 | 0.287 | -0.03 | 0.09 | 643 |

<u>Desk rejections</u>

**SI Table 13 Question: "For each of the journals listed in the table below, please indicate the appropriate number of manuscripts rejected" – mean desk rejection rate**

| | Women | | | | | Men | | | | |
|---|---|---|---|---|---|---|---|---|---|---|
| Desk Rejection Rate of S.N.P. | Obs | Mean | Std. dev. | Min | Max | Obs | Mean | Std. dev. | Min | Max |
| MS | 262 | 0.76 | 0.34 | 0 | 1 | 565 | 0.79 | 0.28 | 0 | 1 |
| NSE | 172 | 0.81 | 0.31 | 0 | 1 | 764 | 0.79 | 0.29 | 0 | 1 |
| SS | 46 | 0.85 | 0.26 | 0 | 1 | 75 | 0.87 | 0.24 | 0 | 1 |
| All disciplines | 480 | 0.79 | 0.32 | 0 | 1 | 1,404 | 0.80 | 0.28 | 0 | 1 |
| | Women | | | | | Men | | | | |
| Desk Rejection Rate of NC.SA. | Obs | Mean | Std. dev. | Min | Max | Obs | Mean | Std. dev. | Min | Max |
| MS | 105 | 0.80 | 0.34 | 0 | 1 | 210 | 0.77 | 0.35 | 0 | 1 |
| NSE | 64 | 0.72 | 0.38 | 0 | 1 | 266 | 0.74 | 0.37 | 0 | 1 |
| SS | 4 | 0.75 | 0.50 | 0 | 1 | 11 | 0.86 | 0.25 | 0 | 1 |

| | | | | | | | | | | |
|---|---|---|---|---|---|---|---|---|---|---|
| All disciplines | 173 | 0.77 | 0.36 | 0 | 1 | 487 | 0.76 | 0.36 | 0 | 1 |
| | Women | | | | | Men | | | | |
| Desk Rejection Rate of NEJM.C. | Obs | Mean | Std. dev. | Min | Max | Obs | Mean | Std. dev. | Min | Max |
| MS | 161 | 0.68 | 0.40 | 0 | 1 | 320 | 0.67 | 0.40 | 0 | 1 |
| NSE | 9 | 0.72 | 0.44 | 0 | 1 | 28 | 0.71 | 0.43 | 0 | 1 |
| SS | 6 | 0.83 | 0.26 | 0.5 | 1 | 13 | 0.88 | 0.30 | 0 | 1 |
| All disciplines | 176 | 0.68 | 0.39 | 0 | 1 | 361 | 0.68 | 0.40 | 0 | 1 |

**SI Table 14 Question: "For each of the journals listed in the table below, please indicate the appropriate number of manuscripts rejected" – linear regression.**

| Journal | Discipline | Coefficient | Std. dev. | t | p>\|t\| | 95% CI Lower | 95% CI Upper | obs |
|---|---|---|---|---|---|---|---|---|
| S.N.P. | MS | -0.03 | 0.02 | -1.33 | 0.185 | -0.07 | 0.01 | 809 |
| | NSE | 0.03 | 0.03 | 1.01 | 0.314 | -0.02 | 0.07 | 928 |
| | SS | 0.00 | 0.05 | -0.09 | 0.928 | -0.09 | 0.09 | 119 |
| | All disciplines | 0.00 | 0.02 | -0.31 | 0.757 | -0.04 | 0.03 | 1,856 |
| NC.SA. | MS | 0.04 | 0.04 | 0.96 | 0.336 | -0.04 | 0.12 | 311 |
| | NSE | -0.03 | 0.05 | -0.61 | 0.542 | -0.14 | 0.07 | 326 |
| | SS | -0.14 | 0.20 | -0.71 | 0.494 | -0.59 | 0.30 | 15 |
| | All disciplines | 0.01 | 0.03 | 0.17 | 0.867 | -0.06 | 0.07 | 652 |

| | | | | | | | | |
|---|---|---|---|---|---|---|---|---|
| NEJM. C. | MS | 0.02 | 0.04 | 0.37 | 0.708 | -0.06 | 0.09 | 467 |
| | NSE | 0.03 | 0.17 | 0.18 | 0.862 | -0.32 | 0.38 | 37 |
| | SS | -0.09 | 0.17 | -0.53 | 0.607 | -0.45 | 0.27 | 19 |
| | All disciplines | 0.01 | 0.04 | 0.33 | 0.741 | -0.06 | 0.09 | 523 |

Reasons for not considering submitting to top journals

**SI Table 15 Question: "Please indicate the reason(s) why you did not consider submitting a manuscript to "journal name taken from response to the previous question" – number of respondents and percentage selecting response by discipline and gender.**

| | MS | | | | NSE | | | | SS | | | |
|---|---|---|---|---|---|---|---|---|---|---|---|---|
| | Women | | Men | | Women | | Women | | Men | | Women | |
| Reasons for not submitting to S.N.P. | Total | %Yes | Total | %Yes | Total | %Yes | Total | %Yes | Total | %Yes | Total | %Yes |
| Work was not of high enough quality | 501 | 36% | 553 | 41% | 328 | 37% | 998 | 27% | 109 | 33% | 110 | 30% |
| Work fell outside the scope of the journal | 501 | 61% | 553 | 55% | 328 | 46% | 998 | 49% | 109 | 72% | 110 | 59% |
| Work was not groundbreaking or sufficiently novel | 501 | 52% | 553 | 50% | 328 | 53% | 998 | 44% | 109 | 53% | 110 | 40% |
| Work would fit better in a more specialized journal | 501 | 56% | 553 | 63% | 328 | 64% | 998 | 70% | 109 | 58% | 110 | 69% |
| Work would reach a wider audience in another journal | 501 | 9% | 553 | 13% | 328 | 16% | 998 | 17% | 109 | 7% | 110 | 8% |
| Co-authors wished to submit the manuscript elsewhere | 501 | 7% | 553 | 7% | 328 | 9% | 998 | 5% | 109 | 6% | 110 | 5% |
| I was advised against submitting to this journal | 501 | 6% | 553 | 4% | 328 | 8% | 998 | 5% | 109 | 6% | 110 | 5% |
| | MS | | | | NSE | | | | SS | | | |
| | Women | | Men | | Women | | Women | | Men | | Women | |
| Reasons for not submitting to NC.SA. | n | %Yes | n | % Yes | n | % Yes | n | % Yes | n | % Yes | n | % Yes |
| Work was not of high enough quality | 667 | 22% | 894 | 23% | 427 | 21% | 1,467 | 15% | 157 | 12% | 180 | 17% |
| Work fell outside the scope of the journal | 667 | 46% | 894 | 37% | 427 | 33% | 1,467 | 34% | 157 | 61% | 180 | 53% |
| Work was not groundbreaking or sufficiently novel | 667 | 32% | 894 | 29% | 427 | 37% | 1,467 | 28% | 157 | 29% | 180 | 23% |
| Work would fit better in a more specialized journal | 667 | 42% | 894 | 47% | 427 | 53% | 1,467 | 52% | 157 | 39% | 180 | 51% |
| Work would reach a wider audience in another journal | 667 | 9% | 894 | 14% | 427 | 13% | 1,467 | 17% | 157 | 10% | 180 | 11% |
| Co-authors wished to submit the manuscript elsewhere | 667 | 9% | 894 | 9% | 427 | 10% | 1,467 | 8% | 157 | 6% | 180 | 4% |
| I was advised against submitting to this journal | 667 | 4% | 894 | 4% | 427 | 3% | 1,467 | 3% | 157 | 3% | 180 | 2% |
| Reasons for not submitting to NEJM.C. | MS | | | | NSE | | | | SS | | | |
| | Women | | Men | | Women | | Women | | Men | | Women | |

| | n | %Yes | n | % Yes | n | % Yes | n | % Yes | n | % Yes | n | % Yes |
|---|---|---|---|---|---|---|---|---|---|---|---|---|
| Work was not of high enough quality | 602 | 34% | 791 | 31% | 503 | 6% | 1,801 | 4% | 151 | 15% | 180 | 8% |
| Work fell outside the scope of the journal | 602 | 83% | 791 | 80% | 503 | 84% | 1,801 | 86% | 151 | 93% | 180 | 90% |
| Work was not groundbreaking or sufficiently novel | 602 | 38% | 791 | 33% | 503 | 8% | 1,801 | 5% | 151 | 21% | 180 | 12% |
| Work would fit better in a more specialized journal | 602 | 37% | 791 | 35% | 503 | 13% | 1,801 | 13% | 151 | 24% | 180 | 27% |
| Work would reach a wider audience in another journal | 602 | 5% | 791 | 7% | 503 | 3% | 1,801 | 4% | 151 | 5% | 180 | 6% |
| Co-authors wished to submit the manuscript elsewhere | 602 | 5% | 791 | 6% | 503 | 2% | 1,801 | 2% | 151 | 5% | 180 | 3% |
| I was advised against submitting to this journal | 602 | 4% | 791 | 4% | 503 | 0% | 1,801 | 0% | 151 | 2% | 180 | 1% |

**SI Table 16 Question: "Please indicate the reason(s) why you did not consider submitting a manuscript to "journal name taken from response to the previous question" – odds ratio (women to men) values.**

| Reasons why did not consider submitting papers to top journals | S.N.P. | | | | NC.SA. | | | | NEJM.C. | | | |
|---|---|---|---|---|---|---|---|---|---|---|---|---|
| | MS | NSE | SS | All displ. | MS | NSE | SS | All displ. | MS | NSE | SS | All displ. |
| Work was not of high enough quality | 0.83 | 1.59** | 1.04 | 1.12 | 0.91 | 1.51** | 0.60 | 1.08 | 1.10 | 1.51 | 1.69 | 1.21 |
| Work fell outside the scope of the journal | 1.27 | 0.94 | 1.84* | 1.14 | 1.42** | 1.01 | 1.41 | 1.24** | 1.20 | 0.92 | 1.41 | 1.08 |
| Work was not groundbreaking or sufficiently novel | 1.24* | 1.47** | 1.69* | 1.26** | 1.13 | 1.41** | 1.27 | 1.26** | 1.18 | 1.60* | 1.71 | 1.31** |
| Work would fit better in a more specialized journal | 0.74* | 0.72* | 0.58 | 0.72* | 0.82 | 1.02 | 0.60* | 0.87 | 1.06 | 1.01 | 0.81 | 1.01 |
| Work would reach a wider audience in another journal | 0.61* | 0.93 | 0.92 | 0.78 | 0.58** | 0.74 | 0.90 | 0.68** | 0.71 | 0.81 | 0.80 | 0.76 |
| Co-authors wished to submit the manuscript elsewhere | 1.02 | 1.70* | 1.31 | 1.32 | 1.00 | 1.17 | 1.34 | 1.09 | 0.88 | 1.09 | 1.34 | 0.97 |
| I was advised against submitting to this journal | 1.30 | 1.69* | 1.37 | 1.48* | 0.87 | 0.77 | 1.73 | 0.87 | 1.24 | 1.11 | 1.52 | 1.25 |
| I was unaware of this journal | + no one reported they were unaware of the top journals | | | | | | | | | | | |

* Indicates p<0.05 ** indicates p<001
+ Insufficient sample

PART 3 Most cited

Journal submission behavior

**SI Table 17 Question: “For your published manuscript <Title of the respondent’s most cited paper>, did you ever consider submitting it to a journal other than <the title of the journal in which it was published>?” – percentage by discipline, and gender.**

| Discipline | Women | | | | Men | | | |
|---|---|---|---|---|---|---|---|---|
| | No | Yes | Total | %Yes | No | Yes | Total | %Yes |
| MS | 370 | 432 | 802 | 53.9% | 559 | 616 | 1,175 | 52.4% |
| NSE | 273 | 248 | 521 | 47.6% | 1,038 | 814 | 1,852 | 44.0% |
| SS | 80 | 84 | 164 | 51.2% | 101 | 92 | 193 | 47.7% |
| All disciplines | 723 | 764 | 1,487 | 51.4% | 1,698 | 1,522 | 3,220 | 47.3% |

**SI Table 18 Question: “For your published manuscript <Title of the respondent’s most cited paper>, did you ever consider submitting it to a journal other than <the title of the journal in which it was published>?” – odds ratio (women to men) values.**

| Other journals | F/M Odds Ratio | Std.err | z | p>\|z\| | 95% CI Lower | 95% CI Upper | obs |
|---|---|---|---|---|---|---|---|
| MS | 1.07 | 0.100 | 0.67 | 0.50 | 0.89 | 1.28 | 1,914 |
| NSE | 1.12 | 0.113 | 1.14 | 0.26 | 0.92 | 1.37 | 2,351 |
| SS | 1.16 | 0.250 | 0.68 | 0.50 | 0.76 | 1.77 | 351 |
| All disciplines | 1.10 | 0.072 | 1.40 | 0.16 | 0.96 | 1.25 | 4,616 |

**SI Table 19 Question: “Before publication of your manuscript, <Title of the respondent’s most cited paper>, did you ever consider submitting it to the journals listed in the table below?” – percentage by discipline, journal, and gender.**

| Consider Submitting to S.N.P. | Women | | | | Men | | | |
|---|---|---|---|---|---|---|---|---|
| | No | Yes | Total | %Yes | No | Yes | Total | %Yes |
| MS | 721 | 76 | 797 | 9.5% | 1,029 | 143 | 1,172 | 12.2% |
| NSE | 478 | 40 | 518 | 7.7% | 1,702 | 148 | 1,850 | 8.0% |
| SS | 157 | 6 | 163 | 3.7% | 185 | 13 | 198 | 6.6% |
| All disciplines | 1,356 | 122 | 1,478 | 8.3% | 2,916 | 304 | 3,220 | 9.4% |
| | Women | | | | Men | | | |
| Consider Submitting to NC.SA. | No | Yes | Total | %Yes | No | Yes | Total | %Yes |
| MS | 743 | 32 | 775 | 4.1% | 1,082 | 46 | 1,128 | 4.1% |
| NSE | 486 | 17 | 503 | 3.4% | 1,747 | 47 | 1,794 | 2.6% |
| SS | 161 | 1 | 162 | 0.6% | 194 | 0 | 194 | 0.0% |

| | | | | | | | | |
|---|---|---|---|---|---|---|---|---|
| All disciplines | 1,390 | 50 | 1,440 | 3.5% | 3,023 | 93 | 3,116 | 3.0% |
| | Women | | | | Men | | | |
| Consider Submitting to NEJM.C. | No | Yes | Total | %Yes | No | Yes | Total | %Yes |
| MS | 749 | 42 | 791 | 5.3% | 1,078 | 73 | 1,151 | 6.3% |
| NSE | 500 | 2 | 502 | 0.4% | 1,788 | 3 | 1,791 | 0.2% |
| SS | 163 | 0 | 163 | 0.0% | 192 | 2 | 194 | 1.0% |
| All disciplines | 1,412 | 44 | 1,456 | 3.0% | 3,058 | 78 | 3,136 | 2.5% |

**SI Table 20 Question: “Before publication of your manuscript, <Title of the respondent’s most cited paper>, did you ever consider submitting it to the journals listed in the table below?” – odds ratio (women to men) values.**

| Journal | | W/M Odds Ratio | Std.err | z | p>\|z\| | 95% CI Lower | 95% CI Upper | obs |
|---|---|---|---|---|---|---|---|---|
| S.N.P. | MS | 0.79 | 0.120 | -1.53 | 0.126 | 0.59 | 1.07 | 1,907 |
| | NSE | 0.96 | 0.181 | -0.21 | 0.83 | 0.66 | 1.39 | 2,342 |
| | SS | 0.54 | 0.275 | -1.21 | 0.227 | 0.20 | 1.47 | 355 |
| | All disciplines | 0.83 | 0.096 | -1.59 | 0.112 | 0.66 | 1.04 | 4,604 |
| NC.SA. | MS | 1.01 | 0.240 | 0.05 | 0.961 | 0.64 | 1.61 | 1,844 |
| | NSE | 1.05 | 0.321 | 0.16 | 0.873 | 0.58 | 1.91 | 2,272 |
| | SS | Insufficient sample | | | | | | |

| | | | | | | | | |
|---|---|---|---|---|---|---|---|---|
| | All disciplines | 1.05 | 0.196 | 0.25 | 0.801 | 0.73 | 1.51 | 4,466 |
| NEJM. C. | MS | 0.84 | 0.170 | -0.87 | 0.386 | 0.56 | 1.25 | 1,882 |
| | NSE | 2.44 | 2.246 | 0.97 | 0.333 | 0.40 | 14.84 | 2,268 |
| | SS | Insufficient sample | | | | | | |
| | All disciplines | 0.85 | 0.166 | -0.85 | 0.393 | 0.57 | 1.24 | 4,501 |

**SI Table 21 Question: "Before publication of your manuscript, <Title of the respondent's most cited paper>, did you ever submit it to the journals listed in the table below?" – percentage by discipline, journal, and gender.**

| | Women | | | | Men | | | |
|---|---|---|---|---|---|---|---|---|
| Submitted to S.N.P. | No | Yes | Total | %Yes | No | Yes | Total | %Yes |
| MS | 37 | 38 | 75 | 50.7% | 58 | 81 | 139 | 58.3% |
| NSE | 22 | 17 | 39 | 43.6% | 64 | 80 | 144 | 55.6% |
| SS | 1 | 4 | 5 | 80.0% | 5 | 8 | 13 | 61.5% |
| All disciplines | 60 | 59 | 119 | 49.6% | 127 | 169 | 296 | 57.1% |
| | Women | | | | Men | | | |
| Submitted to NC.SA. | No | Yes | Total | %Yes | No | Yes | Total | %Yes |
| MS | 19 | 14 | 33 | 42.4% | 25 | 27 | 52 | 51.9% |
| NSE | 13 | 7 | 20 | 35.0% | 35 | 21 | 56 | 37.5% |
| SS | 1 | 0 | 1 | 0.0% | 0 | 0 | 0 | 0 |
| All disciplines | 33 | 21 | 54 | 38.9% | 60 | 48 | 108 | 44.4% |
| | Women | | | | Men | | | |
| Submitted to NEJM.C. | No | Yes | Total | %Yes | No | Yes | Total | %Yes |
| MS | 29 | 13 | 42 | 31.0% | 40 | 31 | 71 | 43.7% |
| NSE | 2 | 0 | 2 | 0.0% | 2 | 0 | 2 | 0.0% |
| SS | 0 | 0 | 0 | 0 | 2 | 0 | 2 | 0.0% |
| All disciplines | 31 | 13 | 44 | 29.5% | 44 | 31 | 75 | 41.3% |

**SI Table 22 Question: "Before publication of your manuscript, <Title of the respondent's most cited paper>, did you ever submit it to the journals listed in the table below?" – odds ratio (women to men) values for the probability of submitting.**

| Journal | Discipline | W/M Odds Ratio | Std.err | z | p>\|z\| | 95% CI Lower | 95% CI Upper | obs |
|---|---|---|---|---|---|---|---|---|
| S.N.P. | MS | 0.75 | 0.22 | -0.99 | 0.321 | 0.43 | 1.32 | 210 |
| | NSE | 0.65 | 0.24 | -1.19 | 0.234 | 0.31 | 1.33 | 180 |
| | SS | 2.50 | 3.14 | 0.73 | 0.465 | 0.21 | 29.25 | 18 |
| | All disciplines | 0.74 | 0.17 | -1.35 | 0.177 | 0.48 | 1.15 | 408 |
| NC.SA. | MS | 0.69 | 0.32 | -0.82 | 0.414 | 0.28 | 1.70 | 84 |
| | NSE | 1.06 | 0.61 | 0.10 | 0.918 | 0.35 | 3.25 | 73 |
| | SS | No one submitted to NC.SA. | | | | | | |
| | All disciplines | 0.80 | 0.29 | -0.61 | 0.540 | 0.40 | 1.62 | 157 |
| NEJM.C. | MS | 0.55 | 0.23 | -1.42 | 0.157 | 0.24 | 1.26 | 111 |
| | NSE | No one submitted to NEJM.C. | | | | | | |
| | SS | No one submitted to NEJM.C. | | | | | | |

| | | | | | | | | |
|---|---|---|---|---|---|---|---|---|
| | All disciplines | 0.57 | 0.24 | -1.35 | 0.178 | 0.25 | 1.29 | 117 |

Desk rejections

**SI Table 23 Question: "Was your manuscript <Title of the respondent's most cited paper> sent out for peer review before being rejected by the journals listed in the table below?" – percentage by discipline, journal, and gender.**

| | Women | | | | Men | | | |
|---|---|---|---|---|---|---|---|---|
| sent out for peer review S.N.P. | No | Yes | Total | %DeskRej | No | Yes | Total | %DeskRej |
| MS | 26 | 10 | 36 | 72.2% | 62 | 19 | 81 | 76.5% |
| NSE | 14 | 3 | 17 | 82.4% | 58 | 22 | 80 | 72.5% |
| SS | 3 | 1 | 4 | 75.0% | 8 | 0 | 8 | 100.0% |
| All disciplines | 43 | 14 | 57 | 75.4% | 128 | 41 | 169 | 75.7% |
| | Women | | | | Men | | | |
| sent out for peer review NC.SA. | No | Yes | Total | %DeskRej | No | Yes | Total | %DeskRej |
| MS | 12 | 2 | 14 | 85.7% | 21 | 7 | 28 | 75.0% |
| NSE | 7 | 1 | 8 | 87.5% | 17 | 4 | 21 | 81.0% |
| SS | No responses | | | | | | | |
| All disciplines | 19 | 3 | 22 | 86.4% | 31 | 11 | 49 | 77.6% |
| | Women | | | | Men | | | |
| sent out for peer review NEJM.C. | No | Yes | Total | %DeskRej | No | Yes | Total | %DeskRej |
| MS | 5 | 8 | 13 | 38.5% | 27 | 5 | 32 | 84.4% |
| NSE | No responses | | | | | | | |
| SS | | | | | | | | |

**SI Table 24 Question: "Was your manuscript <Title of the respondent's most cited paper> sent out for peer review before being rejected by the journals listed in the table below?" – odds ratio (women to men) values.**

| Journal | Discipline | W/M Odds Ratio | Std.err | z | p>\|z\| | 95% CI Lower | 95% CI Upper | obs |
|---|---|---|---|---|---|---|---|---|
| S.N.P. | MS | 0.48 | 0.66 | 0.74 | 0.46 | 0.56 | 3.52 | 114 |

| | | | | | | | | |
|---|---|---|---|---|---|---|---|---|
| | NSE | 1.41 | 0.33 | -1.05 | 0.29 | 0.12 | 1.88 | 95 |
| | SS | Insufficient sample | | | | | | |
| | All disciplines | 1.13 | 0.41 | 0.32 | 0.75 | 0.55 | 2.31 | 221 |
| NC.SA. | MS | 0.43 | 0.38 | -0.95 | 0.34 | 0.07 | 2.46 | 36 |
| | NSE | 0.70 | 0.90 | -0.28 | 0.78 | 0.06 | 8.82 | 27 |
| | SS | No responses | | | | | | |
| | All disciplines | 0.50 | 0.37 | -0.95 | 0.35 | 0.12 | 2.10 | 63 |
| NEJM.C. | MS | 13.07 | 10.96 | 3.07 | 0.00 | 2.53 | 67.60 | 44 |
| | NSE | No responses | | | | | | |
| | SS | | | | | | | |

<u>Reasons for not considering submitting to top journals</u>

**SI Table 25 Question: “Please indicate the reason(s) why you did not consider submitting your manuscript <Title of the respondent’s most cited paper> to the journals listed below.” – number of respondents and percentage selecting response by discipline and gender.**

| | MS | | | | NSE | | | | SS | | | |
|---|---|---|---|---|---|---|---|---|---|---|---|---|
| | Women | | Men | | Women | | Men | | Women | | Men | |
| Reasons for not submitting to S.N.P. | n | %Yes | n | % Yes | n | % Yes | n | % Yes | n | % Yes | n | % Yes |
| Work was not of high enough quality | 719 | 25% | 1,027 | 30% | 475 | 21% | 1,696 | 18% | 156 | 27% | 185 | 18% |
| Work fell outside the scope of the journal | 719 | 45% | 1,027 | 44% | 475 | 35% | 1,696 | 40% | 156 | 62% | 185 | 60% |
| Work was not groundbreaking or sufficiently novel | 719 | 45% | 1,027 | 44% | 475 | 48% | 1,696 | 39% | 156 | 39% | 185 | 30% |

| | | | | | | | | | | | | |
|---|---|---|---|---|---|---|---|---|---|---|---|---|
| Work would fit better in a more specialized journal | 719 | 53% | 1,027 | 53% | 475 | 60% | 1,696 | 63% | 156 | 46% | 185 | 57% |
| Work would reach a wider audience in another journal | 719 | 8% | 1,027 | 9% | 475 | 12% | 1,697 | 14% | 156 | 8% | 185 | 11% |
| Co-authors wished to submit the manuscript elsewhere | 719 | 7% | 1,027 | 6% | 475 | 8% | 1,697 | 4% | 156 | 6% | 185 | 3% |
| I was advised against submitting to this journal | 719 | 5% | 1,027 | 3% | 475 | 3% | 1,696 | 2% | 156 | 2% | 185 | 2% |
| | MS | | | | NSE | | | | SS | | | |
| | Women | | Men | | Women | | Women | | Men | | Women | |
| Reasons for not submitting to NC.SA. | n | %Yes | n | % Yes | n | % Yes | n | % Yes | n | % Yes | n | % Yes |
| Work was not of high enough quality | 732 | 19% | 1,076 | 22% | 481 | 14% | 1,731 | 13% | 159 | 14% | 189 | 12% |
| Work fell outside the scope of the journal | 732 | 41% | 1,076 | 38% | 481 | 31% | 1,731 | 33% | 159 | 63% | 189 | 58% |
| Work was not groundbreaking or sufficiently novel | 732 | 34% | 1,076 | 32% | 481 | 38% | 1,731 | 30% | 159 | 29% | 189 | 21% |
| Work would fit better in a more specialized journal | 732 | 46% | 1,076 | 45% | 481 | 52% | 1,731 | 55% | 159 | 32% | 189 | 43% |
| Work would reach a wider audience in another journal | 732 | 8% | 1,076 | 11% | 481 | 10% | 1,731 | 12% | 159 | 5% | 189 | 7% |
| Co-authors wished to submit the manuscript elsewhere | 732 | 6% | 1,076 | 6% | 481 | 6% | 1,732 | 5% | 159 | 8% | 189 | 2% |
| I was advised against submitting to this journal | 732 | 2% | 1,076 | 2% | 481 | 2% | 1,731 | 2% | 159 | 1% | 189 | 1% |
| | MS | | | | NSE | | | | SS | | | |
| | Women | | Men | | Women | | Women | | Men | | Women | |
| Reasons for not submitting to NEJM.C. | n | %Yes | n | % Yes | n | % Yes | n | % Yes | n | % Yes | n | % Yes |
| Work was not of high enough quality | 738 | 25% | 1,073 | 28% | 495 | 5% | 1,771 | 5% | 161 | 15% | 187 | 7% |
| Work fell outside the scope of the journal | 738 | 72% | 1,073 | 75% | 495 | 85% | 1,772 | 86% | 161 | 91% | 187 | 86% |
| Work was not groundbreaking or sufficiently novel | 738 | 37% | 1,073 | 36% | 495 | 9% | 1,771 | 7% | 161 | 18% | 187 | 11% |
| Work would fit better in a more specialized journal | 738 | 44% | 1,073 | 42% | 495 | 16% | 1,771 | 18% | 161 | 25% | 187 | 30% |
| Work would reach a wider audience in another journal | 738 | 6% | 1,073 | 6% | 495 | 4% | 1,771 | 4% | 161 | 4% | 187 | 7% |

| | | | | | | | | | | | | |
|---|---|---|---|---|---|---|---|---|---|---|---|---|
| Co-authors wished to submit the manuscript elsewhere | 738 | 5% | 1,073 | 4% | 495 | 2% | 1,771 | 1% | 161 | 5% | 187 | 2% |
| I was advised against submitting to this journal | 738 | 2% | 1,073 | 2% | 495 | 0% | 1,771 | 0% | 161 | 1% | 187 | 0% |

**SI Table 26 Question: “Please indicate the reason(s) why you did not consider submitting your manuscript <Title of the respondent’s most cited paper> to the journals listed below.” – odds ratio (women to men) values.**

| Reasons why did not consider submitting the most cited papers to top journals | S.N.P. | | | | NC.SA. | | | | NEJM.C. | | | |
|---|---|---|---|---|---|---|---|---|---|---|---|---|
| | MS | NSE | SS | All | MS | NSE | SS | All | MS | NSE | SS | All |
| Work was not of high enough quality | 0.75* | 1.17 | 1.63 | 0.96 | 0.86 | 1.00 | 1.31 | 0.93 | 0.83 | 0.94 | 2.36* | 0.91 |
| Work fell outside the scope of the journal | 1.05 | 0.82 | 1.15 | 0.95 | 1.09 | 0.93 | 1.30 | 1.04 | 0.86 | 0.94 | 1.52 | 0.92 |
| Work was not groundbreaking or sufficiently novel | 1.02 | 1.39** | 1.48 | 1.19* | 1.05 | 1.28* | 1.81 | 1.14 | 1.05 | 1.28 | 1.81 | 1.14 |
| Work would fit better in a more specialized journal | 0.99 | 0.88 | 0.66 | 0.90 | 1.00 | 0.90 | 0.64 | 0.92 | 1.08 | 0.92 | 0.79 | 1.00 |
| Work would reach a wider audience in another journal | 0.96 | 0.87 | 0.71 | 0.89 | 0.72 | 0.87 | 0.76 | 0.79 | 0.98 | 0.85 | 0.55 | 0.88 |
| Co-authors wished to submit the manuscript elsewhere | 1.13 | 1.57* | 2.26 | 1.35* | 0.99 | 1.06 | 4.53* | 1.12 | 1.14 | 1.41 | 2.07 | 1.26 |
| I was advised against submitting to this journal | 1.75* | 1.51 | 0.97 | 1.59* | 0.96 | 1.06 | 2.27 | 1.03 | 1.47 | 3.52 | + | 1.67 |
| I was unaware of this journal | +no one reported they were unaware of the top journals | | | | | | | | | | | |

* Indicates p<0.05
+ Insufficient sample

PART 4 – Perception of the quality of research

**SI Table 27 Question: “Compared to my peers, I feel that the quality of my research is” – average rank and count by discipline and gender.**

| Women | | | | | | |
|---|---|---|---|---|---|---|
| Discipline | Average Quality Rank | Std. dev | #Good & Excellent | #Average | #Fair & Poor | Total |
| MS | 4.2 | 0.69 | 574 | 74 | 8 | 656 |
| NSE | 4.0 | 0.67 | 352 | 60 | 8 | 420 |
| SS | 4.1 | 0.62 | 135 | 18 | 0 | 153 |

| | | | | | | |
|---|---|---|---|---|---|---|
| All disciplines | 4.1 | 0.68 | 1,061 | 152 | 16 | 1,229 |
| Men | | | | | | |
| Discipline | Average Quality Rank | Std. dev | #Good & Excellent | #Average | #Fair & Poor | Total |
| MS | 4.1 | 0.7 | 841 | 126 | 7 | 974 |
| NSE | 4.2 | 0.6 | 1,363 | 185 | 9 | 1,557 |
| SS | 4.1 | 0.8 | 151 | 25 | 6 | 182 |
| All disciplines | 4.1 | 0.7 | 2,355 | 336 | 22 | 2,713 |

Note: Excellent (5) – Poor (1)

**SI Table 28 Question: “Compared to my peers, I feel that the quality of my research is” – ordinal logistic (women to men), controlled for rank.**

| Disciplines | Odds Ratio | Std. Err | Z | p>\|z\| | 95% CI Lower | 95% CI Upper | obs |
|---|---|---|---|---|---|---|---|
| MS | 1.18 | 0.13 | 2.40 | 0.07 | 0.99 | 1.40 | 1,630 |
| NSE | 0.83 | 0.09 | -1.64 | 0.04 | 0.67 | 0.99 | 1,977 |
| SS | 1.13 | 0.24 | 0.56 | 0.58 | 0.74 | 1.71 | 335 |
| All disciplines | 1.06 | 0.07 | 0.87 | 0.38 | 0.93 | 1.22 | 3,942 |
